# The FDA- approved gold drug Auranofin inhibits novel coronavirus (SARS-COV-2) replication and attenuates inflammation in human cells

**DOI:** 10.1101/2020.04.14.041228

**Authors:** Hussin A. Rothan, Shannon Stone, Janhavi Natekar, Pratima Kumari, Komal Arora, Mukesh Kumar

## Abstract

SARS-COV-2 has recently emerged as a new public health threat. Herein, we report that the FDA-approved gold drug, auranofin, inhibits SARS-COV-2 replication in human cells at low micro molar concentration. Treatment of cells with auranofin resulted in a 95% reduction in the viral RNA at 48 hours after infection. Auranofin treatment dramatically reduced the expression of SARS-COV-2-induced cytokines in human cells. These data indicate that auranofin could be a useful drug to limit SARS-CoV-2 infection and associated lung injury due to its anti-viral, anti-inflammatory and anti-ROS properties. Auranofin has a well-known toxicity profile and is considered safe for human use.

---

Gold-based compounds have shown promising activity against a wide range of clinical conditions and microorganism infections. Auranofin, a gold-containing triethyl phosphine, is an FDA-approved drug for the treatment of rheumatoid arthritis since 1985 (1). It has been investigated for potential therapeutic application in a number of other diseases including cancer, neurodegenerative disorders, HIV/AIDS, parasitic infections and bacterial infections (1, 2). Recently, auranofin was approved by FDA for phase II clinical trials for cancer therapy. The mechanism of action of auranofin involves the inhibition of redox enzymes such as thioredoxin reductase, induction of endoplasmic reticulum (ER) stress and subsequent activation of the unfolded protein response (UPR) (2–5). Inhibition of these redox enzymes leads to cellular oxidative stress and intrinsic apoptosis (6, 7). In addition, auranofin is an anti-inflammatory drug that reduces cytokines production and stimulate cell-mediated immunity (8). The dual inhibition of inflammatory pathways and thiol redox enzymes by auranofin makes it an attractive candidate for cancer therapy and treating microbial infections.

Coronaviruses are a family of enveloped viruses with positive sense, single-stranded RNA genomes (9). SARS-CoV-2, the causative agent of COVID-19, is closely related to severe acute respiratory syndrome coronavirus (SARS-CoV-1) (9, 10). It is known that ER stress and UPR activation contribute significantly to the viral replication and pathogenesis during a coronavirus infection (11). Infection with SARS-COV-1 increases the expression of the ER protein folding chaperons GRP78, GRP94 and other ER stress related genes to maintain protein folding (12). Cells overexpressing the SARS-COV spike protein and other viral proteins exhibit high levels of UPR activation (13, 14). Thus, inhibition of redox enzymes such as thioredoxin reductase and induction of ER stress by auranofin could significantly affect SARS-COV-2 protein synthesis (15).

In addition, SARS-COV-2 infection causes acute inflammation and neutrophilia that leads to a cytokine storm with over expression of TNF-alpha, monocyte chemoattractant protein (MCP-1) and reactive oxygen species (ROS) (10). The severe COVID-19 illness represents a devastating inflammatory lung disorder due to cytokines storm that is associated with multiple organ dysfunction leading to high mortality (10, 16). Taken together, these studies suggest that auranofin could mitigate SARS-COV-2 infection and associated lung damage due to its anti-viral, anti-inflammatory and anti-ROS properties. Auranofin has a well-known toxicity profile and is considered safe for human use.

We investigated the anti-viral activity of auranofin against SARS-CoV-2 and its effect on virus-induced inflammation in human cells. We infected Huh7 cells with SARS-CoV-2 (USA-WA1/2020) at a multiplicity of infection (MOI) of 1 for 2 hours, followed by the addition of 4 μM of auranofin (17, 18). DMSO (0.1%) was used as control (the solvent was used to prepare drug stock). Cell culture supernatants and cell lysates were collected at 24 and 48 hours after infection. Virus RNA copies were measured by RT-PCR using two separate primers specific for the viral N1 gene and N2 gene (19, 20). As depicted in Figure 1, treatment of cells with auranofin resulted in a 70% reduction in the viral RNA in the supernatant compared to the DMSO at 24 hours after infection. At 48 hours, there was an 85% reduction in the viral RNA in the supernatant compared to the DMSO. Similarly, the levels of intracellular viral RNA decreased by 85% at 24 hours and 95% at 48 hours in auranofin-treated cells compared to the DMSO-treated cells. Both set of primers showed nearly identical results. Auranofin showed no toxicity against Huh7 cells at the used concentration at 24 and 48 hours.

**Figure 1:**
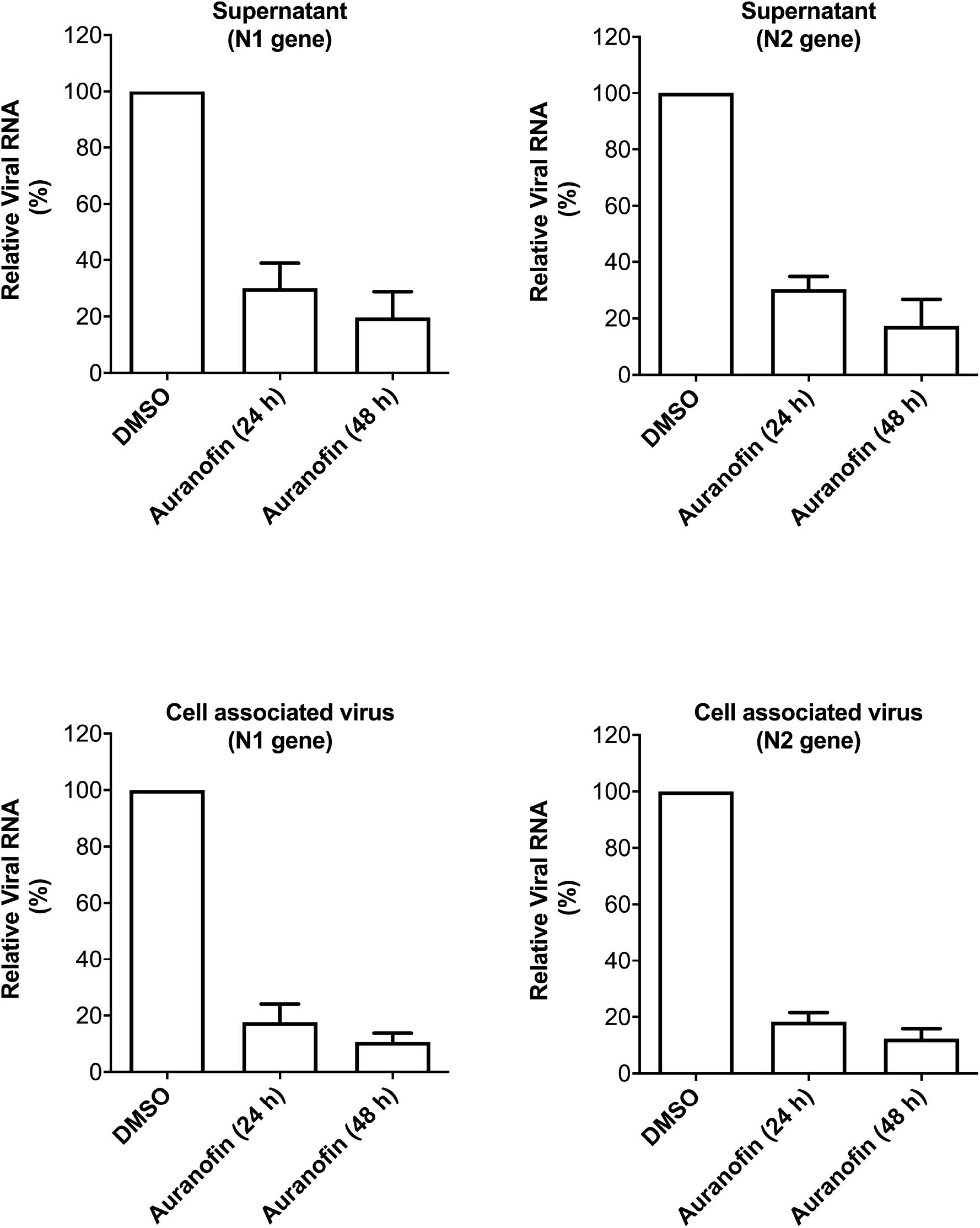
Auranofin inhibits replication of SARS-COV-2 in human cells. Huh7 cells were infected with SARS-COV-2 at a multiplicity of infection (MOI) of 1 for 2 hours and treated with 4 μM of auranofin or with 0.1% DMSO. Cell pellets and culture supernatants were collected at 24 and 48 hours after infection and viral RNA levels were measured by RT-PCR using primers and probe targeting the SARS-COV-2 N1 gene and the SARS-COV-2 N2 gene. The results were identical for both set of primers showing dramatic reduction in viral RNA at both 24 and 48 hours. Data represent the mean±SEM, representing two independent experiments conducted in duplicate.

To determine the effective concentration of auranofin that inhibits 50% of viral replication (EC_50_), we treated SARS-COV-2 infected Huh7 cells with serial dilutions of auranofin. Supernatants and cell lysates were collected at 48 hours after infection and viral RNA was quantified by RT-PCR. The data were plotted in graphs using non-linear regression model (GraphPad software). At 48 hours, there was a dose-dependent reduction in viral RNA levels in the auranofin-treated cells. Figure 2 represents the EC_50_ values of auranofin treatment against SARS-CoV-2 infected Huh7 cells. Auranofin inhibited virus replication in the infected cells at EC_50_ of approximately 1.5 μM.

**Figure 2:**
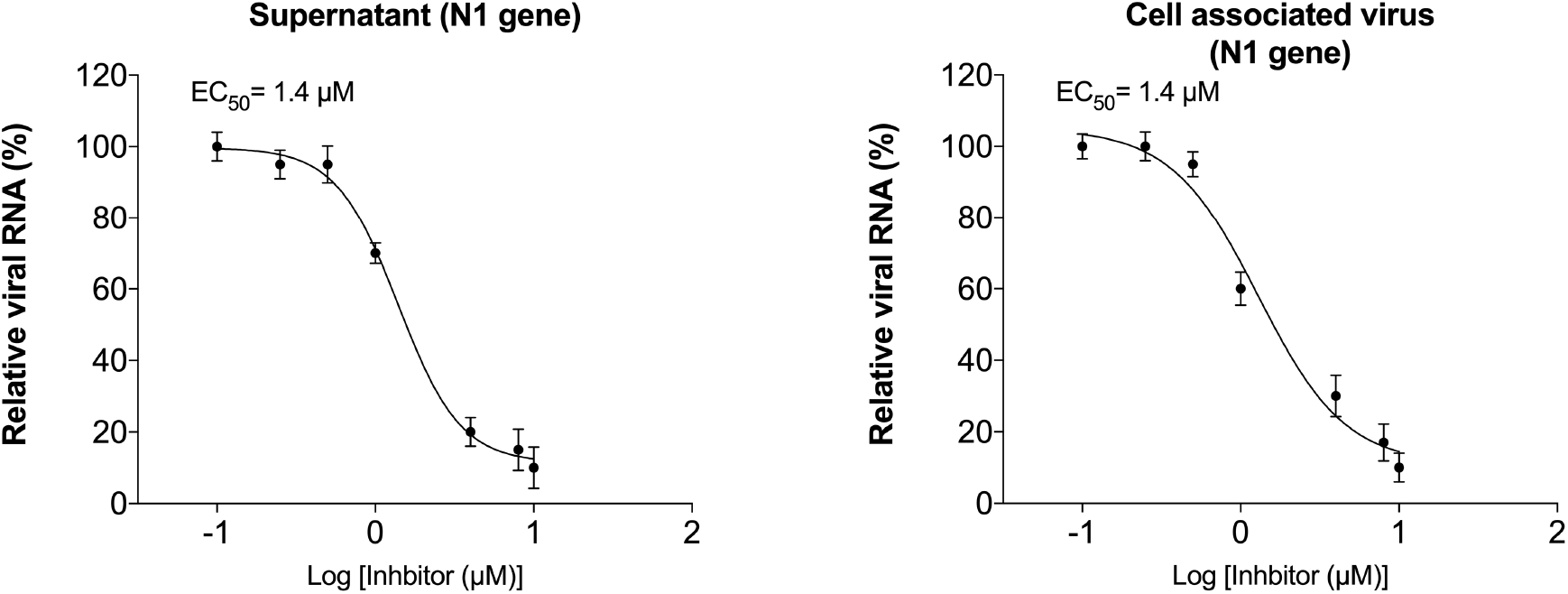
Dose-dependent reduction in SARS-COV-2 RNA in the auranofin-treated cells: The SARS-COV-2 infected Huh7 cells were treated with serial dilutions of auranofin. Viral RNA in the cell pellets and culture supernatants were quantified by RT-PCR using primers and probe targeting the SARS-COV-2 N1. The data were plotted in graphs using non-linear regression model (GraphPad software). Auranofin inhibited virus replication in the infected cells at EC_50_ of approximately 1.5 μM. Data represent two independent experiments conducted in duplicate.

To assess the effect of auranofin on inflammatory response during SARS-COV-2 infection, we measured the levels of key cytokines in auranofin and DMSO-treated cells at 24 and 48 hours after infection (21). SARS-COV-2 infection induces a strong up-regulation of IL-6, IL-1β, TNFα and NF-kB in Huh7 cells (Figure 3). Treatment with auranofin dramatically reduced the expression of SARS-COV-2-induced cytokines in Huh7 cells. SARS-COV-2 infection resulted in a 200-fold increase in the mRNA expression of IL-6 at 48 hours after infection compared to corresponding mock-infected cells. In contrast, there was only a 2-fold increase in expression of IL-6 in auranofin-treated cells. TNF-α levels increased by 90-fold in the DMSO-treated cells at 48 hours after infection, but this increase was absent in the auranofin-treated cells. Similarly, no increase in the expression of IL-1β and NF-kB was observed in the auranofin-treated cells.

**Figure 3:**
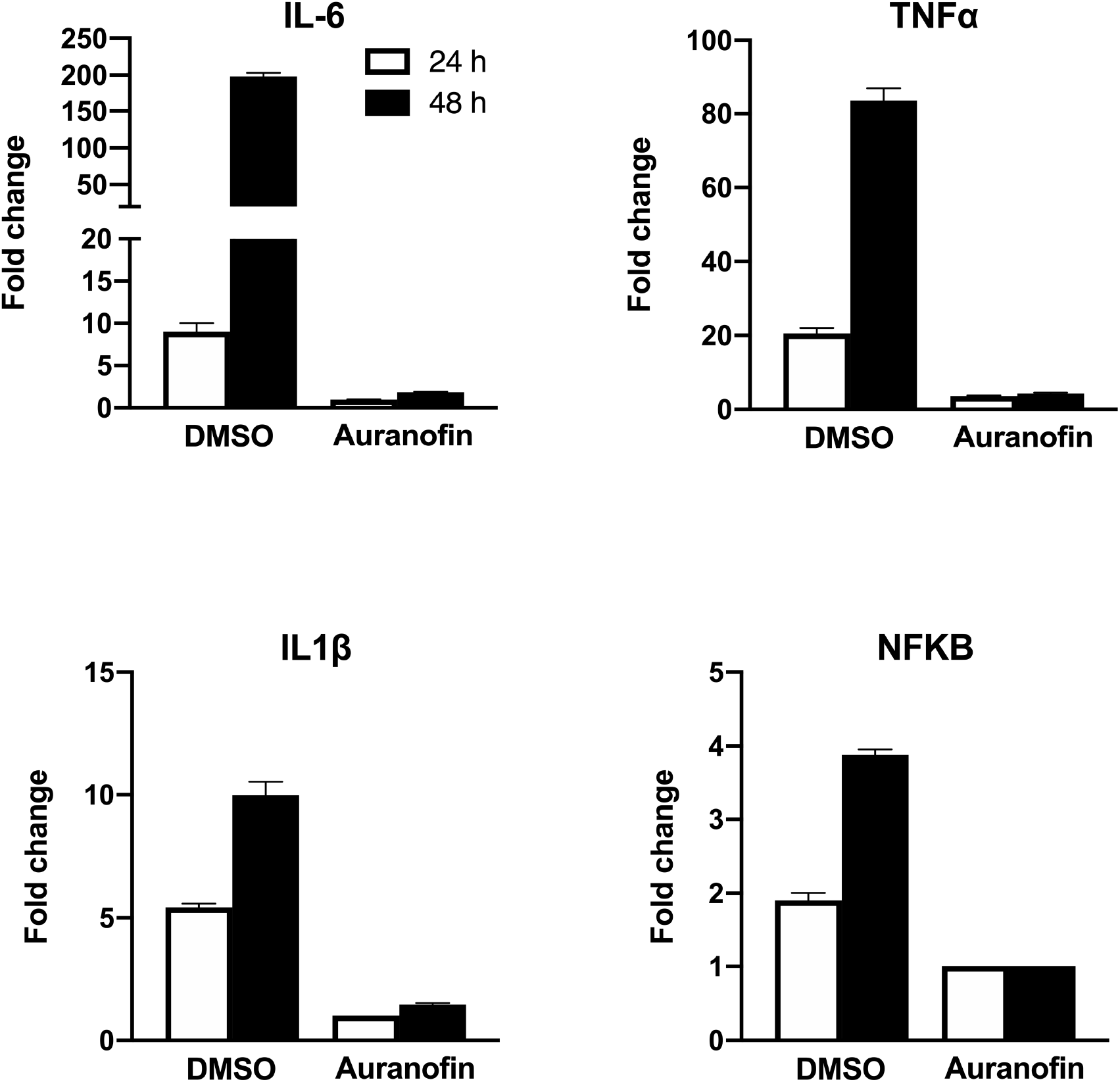
Auranofin treatment dramatically reduced the expression of SARS-COV-2-induced cytokines in human cells: mRNA levels of IL-6, IL-1β, TNFα and NF-kB were determined using qRT-PCR at 24 and 48 hours after infection. The fold change in infected cells compared to corresponding controls was calculated after normalizing to the GAPDH gene. Data represent the mean±SEM, representing two independent experiments conducted in duplicate.

**Taken together** these results demonstrate that auranofin inhibits replication of SARS-COV-2 in human cells at low micro molar concentration. We also demonstrate that auranofin treatment resulted in significant reduction in virus-induced inflammation. These data indicate that auranofin could be a useful drug to limit SARS-CoV-2 infection and associated lung injury. Further animal studies are warranted to evaluate the efficacy of auranofin for the management of SARS-COV-2 associated disease.

## Methods

### SARS-COV-2 infection and drug treatment

In this study, we used a novel SARS-COV-2 (USA-WA1/2020) isolated from an oropharyngeal swab from a patient in Washington, USA (BEI NR-52281). Virus strain was amplified once in Vero E6 cells and had titers of 5 × 10^6^ plaque-forming units (PFU)/mL. Huh7 cells (human liver cell line) were grown in DMEM (Gibco) supplemented with 5% heat-inactivated fetal bovine serum. Cells were infected with SARS-COV-2 or PBS (Mock) at a multiplicity of infection (MOI) of 1 for 2 hours (17, 18, 21, 22). Cell were washed twice with PBS and media containing different concentrations of auranofin (Sigma) or DMSO (Sigma) was added to cells. Supernatants and cell lysates were harvested at 24 and 48 hours after infection.

### Viral RNA quantification

Virus RNA levels were analyzed in the supernatant and cell lysates by quantitative reverse transcription-polymerase chain reaction (qRT-PCR). RNA from cell culture supernatants was extracted using a Viral RNA Mini Kit (Qiagen) and RNA from cell lysates was extracted using a RNeasy Mini Kit (Qiagen) as described previously (20, 21, 23). qRT-PCR was used to measure viral RNA levels using previously published primers and probes specific for the SARS-COV-2. Forward (5′-GACCCCAAAATC AGCGAAAT-3′), reverse (5′-TCTGGTTACTGCCAGTTGAATCTG-3′), probe, (5′-FAM-ACCCCGCATTACGTTTGGTGGACC-BHQ1-3’) targeting the SARS-COV-2 N1 gene and Forward (5′-TTACAAACATTGGCCGCAAA-3′), reverse (5′-GCGCGACATTCCGAAGAA3′), probe, (5′-FAM-ACAATTTGCCCCCAGCGCTTCAG-BHQ1-3’) targeting the SARS-COV-2 N2 gene (Integrated DNA Technologies). Viral RNA copies were determined after comparison with a standard curve produced using serial 10-fold dilutions of SARS-COV-2 RNA (18, 20).

### Cytokine analysis

For mRNA analysis of IL-6, IL-1β, TNFα and NF-kB, cDNA was prepared from RNA isolated from the cell lysates using a iScript™ cDNA Synthesis Kit (Bio-Rad, Hercules, CA, USA), and qRT-PCR was conducted as described previously (21, 23, 24). The primer sequences used for qRT-PCR are listed in Table 1.

**Table 1.**
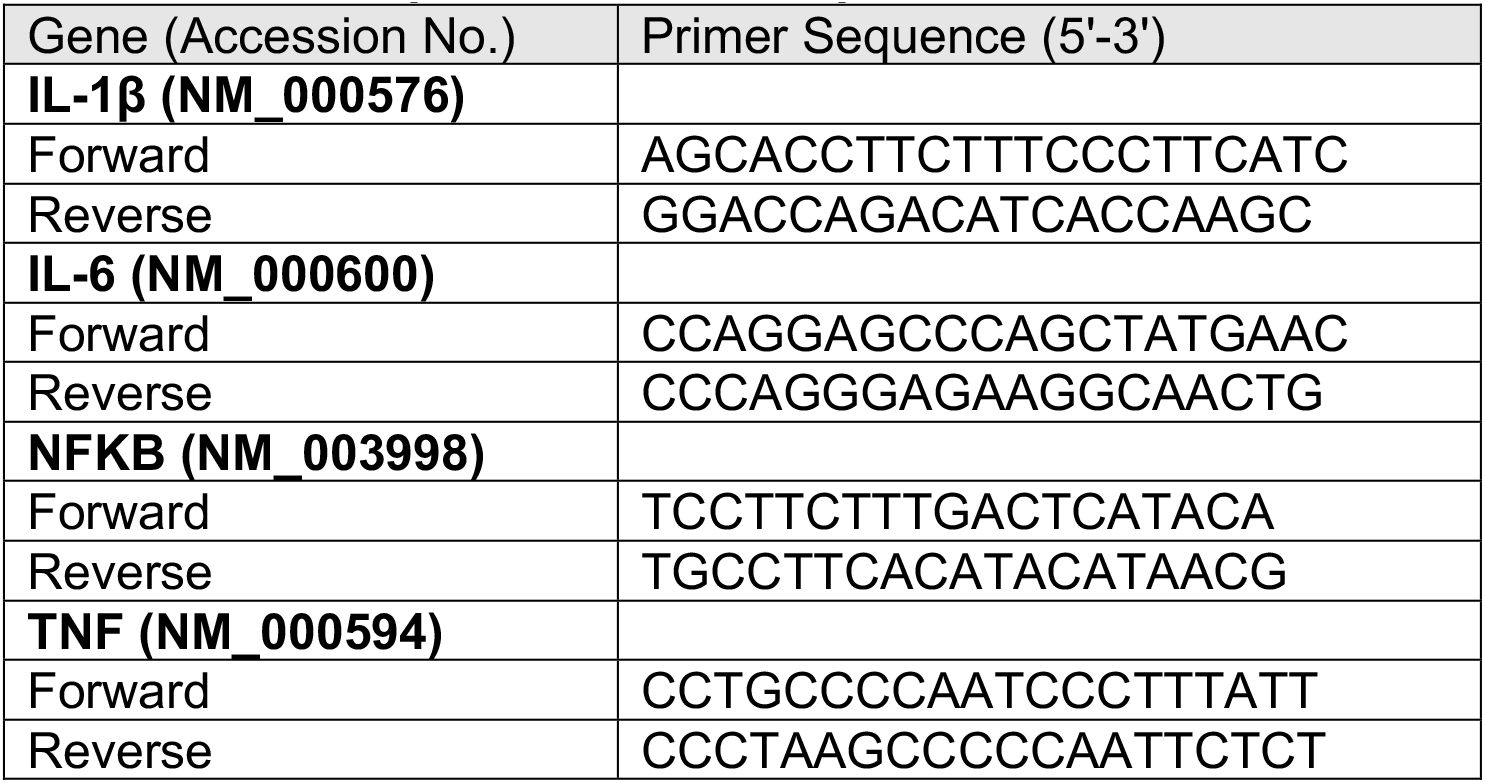
Primer sequences used for qRT-PCR

## Acknowledgements

This work was supported by a grant (R21NS099838) from National Institute of Neurological Disorders and Stroke, grant (R21OD024896) from the Office of the Director, National Institutes of Health, and Institutional funds.

